# Mannose is crucial for mesoderm specification and symmetry breaking in gastruloids

**DOI:** 10.1101/2023.06.05.543730

**Authors:** Chaitanya Dingare, Jenny Yang, Ben Steventon

## Abstract

Patterning and growth are fundamental features of embryonic development that must be tightly coordinated during morphogenesis. As metabolism can control cell growth while also providing mechanistic links to developmental signalling pathways, it is ideally placed to enable this coordination. To understand how metabolism impacts early mesoderm specification, we used mouse embryonic stem (ES) cell-derived gastruloids, as these enable temporal control over metabolic manipulations and can be generated in large quantities. Gastruloids show mosaic expression of two glucose transporters, *Slc2a1* and *Slc2a3* both of which co-express with the expression of both the mesodermal marker *T/Bra* and the neural marker *Sox2*. To understand the significance of cellular glucose uptake in development, we used the glucose metabolism inhibitor 2-deoxy-D-glucose (2-DG). 2-DG specifically blocks the expression of *T/Bra* without affecting the expression of *Sox2* and abolishes axial elongation in gastruloids. Surprisingly, removing glucose completely from the medium did not phenocopy 2-DG treatment despite a significant decline in glycolytic intermediates occurring under both conditions. As 2-DG can also act as a competitive inhibitor of mannose, we added mannose together with 2-DG and found that it could rescue the mesoderm specification. Together, our results show that while mannose is crucial for mesoderm specification, the glycolytic pathway is dispensable at early stages of *T/Bra* expression in gastruloids.

## Introduction

As a tissue or an organ is formed during embryonic development, a series of specification events generate a wide array of different cell types. At the same time, these different cell populations grow at specific rates. These two processes must be tightly coordinated to yield well-proportioned functional tissues and organs. Pre-implantation mouse embryos rely on pyruvate as a carbon source (Nagaraj et al., 2017). However, as gastrulation begins, peri- and post-implantation embryos rely more on glucose, concomitant with an acceleration in embryo size from 600 cells to 90000 cells in just two days (Chi et al., 2020; Clough & Whittingham, 1983; Kelly & West, 1996; Kojima et al., 2014). This raises a key question whether glucose metabolism plays an important role during gastrulation to bring about the coordination among growth, patterning, and morphogenesis.

Glucose metabolism comprises a complex set of various metabolic pathways including various catabolic reactions (producing ATP, NADPH, NADH) and anabolic reactions (nucleotide, amino acid, glycogen synthesis). Additionally, glucose is converted into other hexoses such as mannose and galactose. These hexoses are further modified to form metabolic intermediates such as N-acetylglucosamine and N-acetyl galactosamine that are used in protein glycosylation, an important post-translational modification that dictates the intracellular localisation of proteins. Therefore, there are multiple ways in which glucose metabolism has the potential to coordinate growth and cell fate specification (Miyazawa & Aulehla, 2018).

Qualitative studies have been performed on chicken embryos to assess how different embryonic parts respond to glucose deprivation. The first tissue to show degeneration upon removal of glucose was the tailbud of the embryo (described as the ‘morphogenetically active’ region), highlighting the importance of glucose metabolism in this tissue (Spratt, 1950). This finding was more recently corroborated - blocking glucose metabolism using a glycolysis inhibitor 2-deoxy-D-glucose (2-DG) affects the tailbud and the axial elongation in the chicken embryo. The tailbud of the mouse and the chicken embryo harbours bipotent neuro-mesodermal progenitor populations (NMPs) which contribute neural and mesodermal cells to the neural tube and the pre-somitic mesoderm (PSM) respectively. 2-DG blocks mesoderm specification in the NMP population of the chicken embryo via inhibition of the Wnt signalling pathway (Oginuma et al., 2017). It was later shown that glycolysis-dependent lactic acid produced in and secreted by the tailbud cell population was responsible for maintaining a basic intracellular pH subsequently regulating the acetylation of Beta-catenin and activation of Wnt signalling (Oginuma et al., 2017, 2020). Another study in the mouse embryo where tailbud explants were incubated in the glucose free medium affected PSM development and subsequently segmentation. It was later shown that glycolytic flux also regulates Wnt signalling and the PSM development in the mouse embryo (Bulusu et al., 2017; Miyazawa et al., 2022). Together, these studies highlight a conserved importance of glucose metabolism in mesoderm development during tailbud stages of development which represent the much later stage of embryonic development (posterior body elongation). However, it is unknown whether glucose metabolism plays an any role in mesoderm development during gastrulation – a key stage when much of the mesoderm is formed, embryonic symmetry is broken, and the anterior-posterior axis is set up.

Mouse mutants for glycolytic enzymes hexokinase II and glucose phosphate isomerase show defects in germ layer specification and die post gastrulation (Heikkinen et al., 1999; Kelly & West, 1996). A similar phenotype was observed in mutants of the glucose transporters Glut1 and Glut3: both have poorly formed germ layers and die post gastrulation (Heilig et al., 2003; Schmidt et al., 2009). To assess the role of glucose metabolism specifically in the earliest steps of mesoderm specification, we made use of gastruloids, aggregates derived from mouse embryonic stem cells (ESCs) (Brink et al., 2014). *Brachyury* (*T/Bra*)-expressing mesodermal cells are specified upon Wnt pathway activation, and subsequently aggregate at one pole marking it as the posterior, breaking symmetry of the gastruloids (Brink et al., 2014; Turner et al., 2017). These specification, patterning and morphogenetic events resemble those in vivo mouse embryos, yet occur in the absence of extra-embryonic tissues and signals (Beccari et al., 2018; Turner et al., 2017). Furthermore, in gastruloids, mesoderm specification and axial elongation are temporally separated, making it possible to perturb metabolic enzymes in time-windows ascribed to each biological process using small molecule inhibitors (López-Anguita et al., 2022). The effect of such treatments can be easily monitored in a high-throughput and non-invasive manner. Therefore, gastruloids offer a simple and high-throughput model system to shed light on the role of glucose metabolism in mesoderm specification.

To address the role of glucose metabolism in mesoderm specification, we first analysed the expression pattern of glucose transporters and found that *Slc2a1* and *Slc2a3* are specifically expressed in the mesodermal cells and neural cells. We further tested the significance of this specific expression pattern of glucose transporters by using 2-DG. 2-DG treatment specifically blocks the specification of mesodermal cells, impairing axial elongation. Even though our metabolomics data suggested that 2-DG efficiently blocked glycolysis in the gastruloids, removing glucose completely from the medium does not affect mesoderm specification and axial elongation. This result highlights an unexplored and glycolysis independent effect of 2-DG on mesoderm specification. Since 2-DG shares a structural similarity with D-Mannose it can become incorporated in place of D-Mannose during glycosylation of different proteins (Ma et al., 2022). We tested the possibility that our 2-DG phenotype results from competitive inhibition of mannose by adding D-Mannose to 2-DG treated gastruloids. Strikingly, this treatment not only rescued the mesoderm specification phenotype at molecular level but also axial elongation. Our study thus highlights a key role for mannose in mesoderm specification during gastrulation.

## Results

### *Slc2a1*/Glut1 and *Slc2a3*/Glut3 become localised to the elongating pole of gastruloids at the onset of symmetry breaking

Cellular glucose uptake is facilitated by a special type of transporter belonging to the solute carrier family 2 (Slc) (Thorens & Mueckler, 2010). To test whether gastruloids express glucose transporters, we analysed publicly available single cell RNA-seq and tomo-seq databases (Brink et al., 2020) and found two genes encoding glucose transporters (*Slc2a1* and *Slc2a3*) are expressed in the posterior region of the gastruloids, which harbours mesodermal and neural progenitor cell populations (Brink et al., 2020). To confirm and further establish the spatial expression of glucose transporters in gastruloids, we performed hybridisation chain reaction (HCR) staining for *Slc2a1* and *Slc2a3* along with mesodermal marker *T/Bra* and neural marker *Sox2* (Choi et al., 2018). We also performed immunostaining for Glut1 and Glut3 encoded by *Slc2a1* and *Slc2a3*, respectively, to corroborate the mRNA expression pattern relative to *T/Bra*. Since *T/Bra* begins to be expressed after Wnt activation, we performed a time series experiment between day 3 and day 5. In day 3 gastruloids at the

mRNA level, *Slc2a1* showed a salt-and-pepper expression pattern relative to *T/Bra* and *Sox2* while *Slc2a3* expressed in the core of the gastruloids (Figure 1A-A’’ and 1D-D’’). At protein level, both Glut1 and Glut3 localised to the membrane and showed uniform localisation across the gastruloid (Figure 1G-G’’ and J-J’’). In day 4 gastruloids, *T/Bra* and *Sox2* expression domains were spatially segregated with a marginal overlap while both *Slc2a1* and *Slc2a3* were co-expressed with *T/Bra* and *Sox2*. (Figure 1B-B’’ and E-E’’). This co-localisation with *T/Bra* is also observed at the protein level (Figure 1H-H’’ and K-K’’). In day 5 gastruloids, when the gastruloids have an elongated morphology, both *Slc2a1* and *Slc2a3* showed expression in the posterior zone coinciding with the expression of *T/Bra* and *Sox2*, and in *Sox2* positive cells spread along the anterior-posterior axis (Figure 1C-C’’ and F-F’’). The localisation pattern of Glut1 and Glut3 protein matches that of mRNA distribution in day 5 gastruloids, with both Glut1 and Glut3 localised to *T/Bra* positive cells (Figure 1I-I’’ and L-L’’). Altogether our data showed that the expression and localisation pattern of *Slc2a1*/Glut1 and *Slc2a3*/Glut3 progressively became restricted to both the mesodermal and neural cells co-incident with gastruloid symmetry breaking. 2-deoxy-D-glucose (2-DG), a competitive inhibitor of hexokinase, inhibits mesoderm specification and gastruloid elongation.

**Figure 1:**
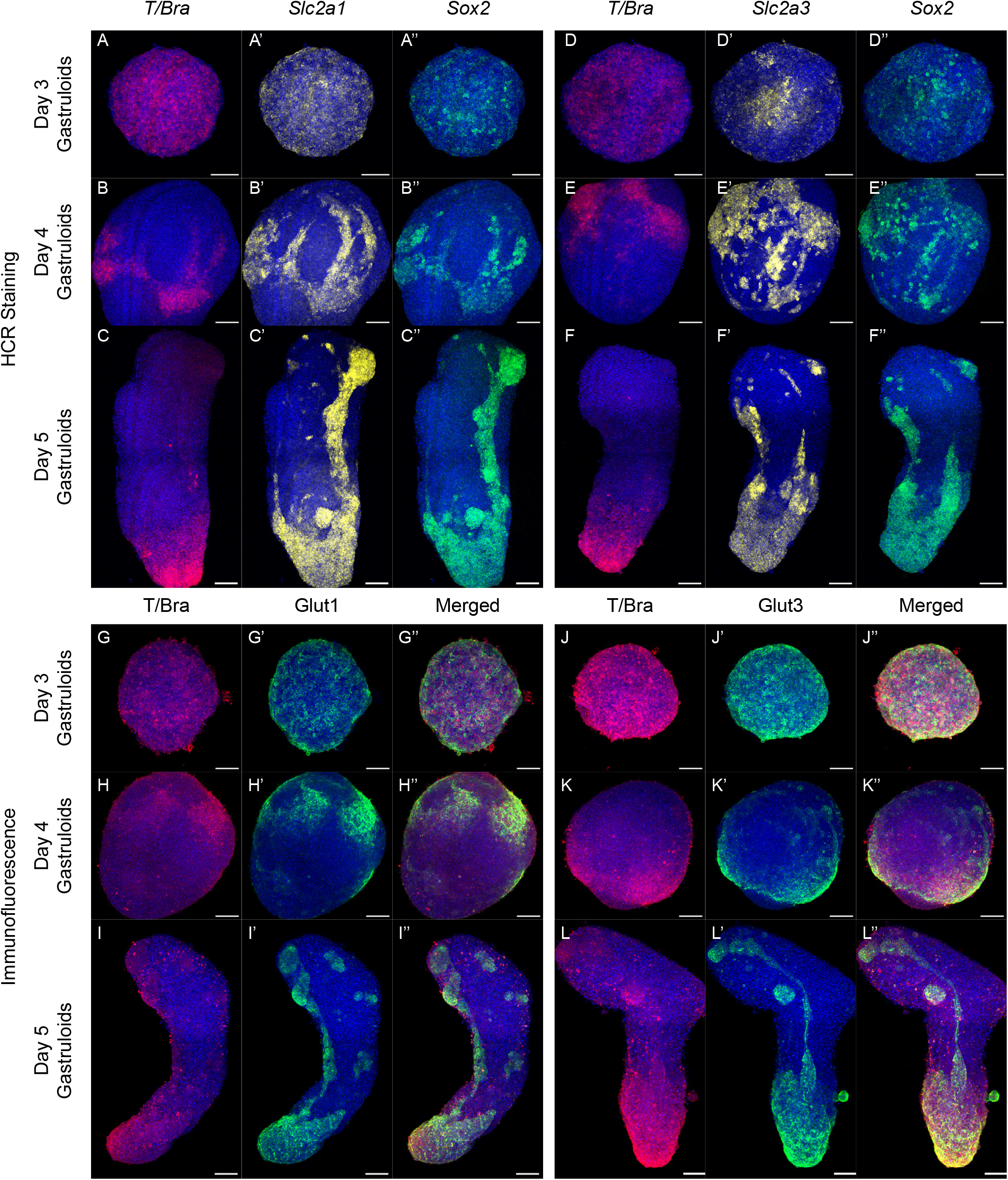
Temporal expression and localisation patterns of Glut1 and Glut3 (A – C’’) Maximum intensity projections of HCR staining showing the expression pattern of *Slc2a1* (Yellow, A’, B’ and C’) with respect to *T/Bra* (Red, A, B, C) and *Sox2* (Green, A’’, B’’ and C’’) in Day 3 (A -A’’, n = 13), Day 4 (B – B’’, n=12) and Day 5 (C – C’’, n=8) gastruloids. (D – F’’) Maximum intensity projections of the HCR staining showing the expression pattern of *Slc2a3* (Yellow, D’, E’ and F’) with respect to *T/Bra* (Red, D, E, F) and *Sox2* (Green, D’’, E’’ and F’’) in Day 3 (D - D’’, n = 13), Day 4 (E – E’’, n=14) and Day 5 (F – F’’, n=8) gastruloids. (G – I’’) Maximum intensity projections of antibody staining showing the localisation pattern of Glut1 (Green, G’, H’ and I’) with respect to *T/Bra* (Red, G, H, I) and co-localisation pattern in merged images (G’’, H’’ and I’’) in Day 3 (G - G’’, n = 15), Day 4 (H – H’’, n=15) and Day 5 (I – I’’, n=10) gastruloids. (J – L’’) Maximum intensity projections of the antibody staining showing the localisation pattern of Glut3 (Green, J’, K’ and L’) with respect to *T/Bra* (Red, J, K, L) and co-localisation pattern in merged images (J’’, K’’ and L’’) in Day 3 (J - J’’, n = 15), Day 4 (K – K’’, n=15) and Day 5 (L – L’’, n=10) gastruloids. Scale bar - 100 μm. The blue staining denotes nuclei in all the confocal images.

Since *Slc2a1* and *Slc2a3* were expressed specifically in mesodermal and neural cells, we wanted to test whether blocking glycolysis affected these cells and subsequently the morphogenesis of the gastruloids. To test this, we treated gastruloids with 2-DG, which blocks the first and second steps of glycolysis, at defined intervals of the gastruloid protocol (Ralser et al., 2008). When treated between days 4 and 5, we observed a minor effect on gastruloid elongation (Figure 2A-A’). The ratio of the major axis that runs from the anterior to the posterior parts of the gastruloids to the minor axis (a proxy for elongation) was mildly reduced in 2-DG treated gastruloids as compared to that of the controls (Figure 2C). In addition, no patterning defects were observed in mesodermal and neural cells except the expression of *T/Bra* appeared to be reduced as compared to the control (Figure 2D-D’). Since glucose transporters were also expressed in earlier stages of gastruloid development, we treated gastruloids with 2-DG between day 3 and 4 (prior to symmetry breaking and elongation). At day 4, gastruloids were transferred to medium containing no 2-DG. Surprisingly, the gastruloids did not elongate, despite experiencing a similar length (24-hour) of treatment with 2-DG as the Day 4-5 treatment (Figure 2B-B’, C). Expression of *T/Bra* could not be observed in the treated gastruloids while the expression of *Sox2* remained unchanged (Figure 2E-E’). To further analyse this phenotype, we performed transcriptomic analysis of day 4 control and 2-DG (5mM) treated gastruloids and computed a list of differentially expressed genes between the two treatments. The gene ontology (GO) term analysis of these genes output terms including anterior/posterior patterning, cell differentiation and embryonic skeletal system development (Figure 2F). Further analysis of the downregulated genes in 2-DG treated gastruloids revealed that many of these genes are related to mesoderm cell fate specification (*T/Bra, Wnt3a, Noto* etc) while the upregulated genes in 2-DG treated gastruloids were related to neural development (*Sox1, Dbx1, Eya1*). We also observed genes related to Wnt signalling (*Lef1*) and to FGF signalling (*Spry4, Etv1*) downregulated in 2-DG treated gastruloids. In addition, many 5’ (posteriorly-expressed) Hox genes were shown to have reduced expression pointing towards the overall suppression of or delay in the posterior fate specifications in the gastruloids upon 2-DG treatment (Figure 2G). To validate the key expression changes in our transcriptomic dataset, we performed HCR staining to observe their expression pattern in control and 2-DG treated gastruloids. Early primitive streak markers i.e., *T/Bra* (Figure 2H -H’), *Wnt3a* (Figure 2I-I’), an FGF target gene *Etv4* (Figure 2L-L’), the Wnt target gene *Lef1* (Figure 2J -J’) and the late Hox gene *Hoxc6* (Figure 2O-O’) all showed a corresponding reduction in the 2-DG treated gastruloids while the expression of the neural marker *Sox2* (Figure 2M-M’) remained unchanged and Sox1 (Figure 2N-N’) showed a mild increase in the treated gastruloids (Yamaguchi et al., 1999). Even though the *Wnt3a* mediated *T/Bra* expression is abrogated upon 2-DG treatment, surprisingly, we did not observe any effect on a broad range Wnt target *Axin2* (Figure 2K-K’), hinting only a part of the Wnt pathway was affected. Altogether our transcriptomics and HCR data showed that 2-DG had an effect at multiple levels i.e., mesoderm specification, posterior development, and signalling.

**Figure 2:**
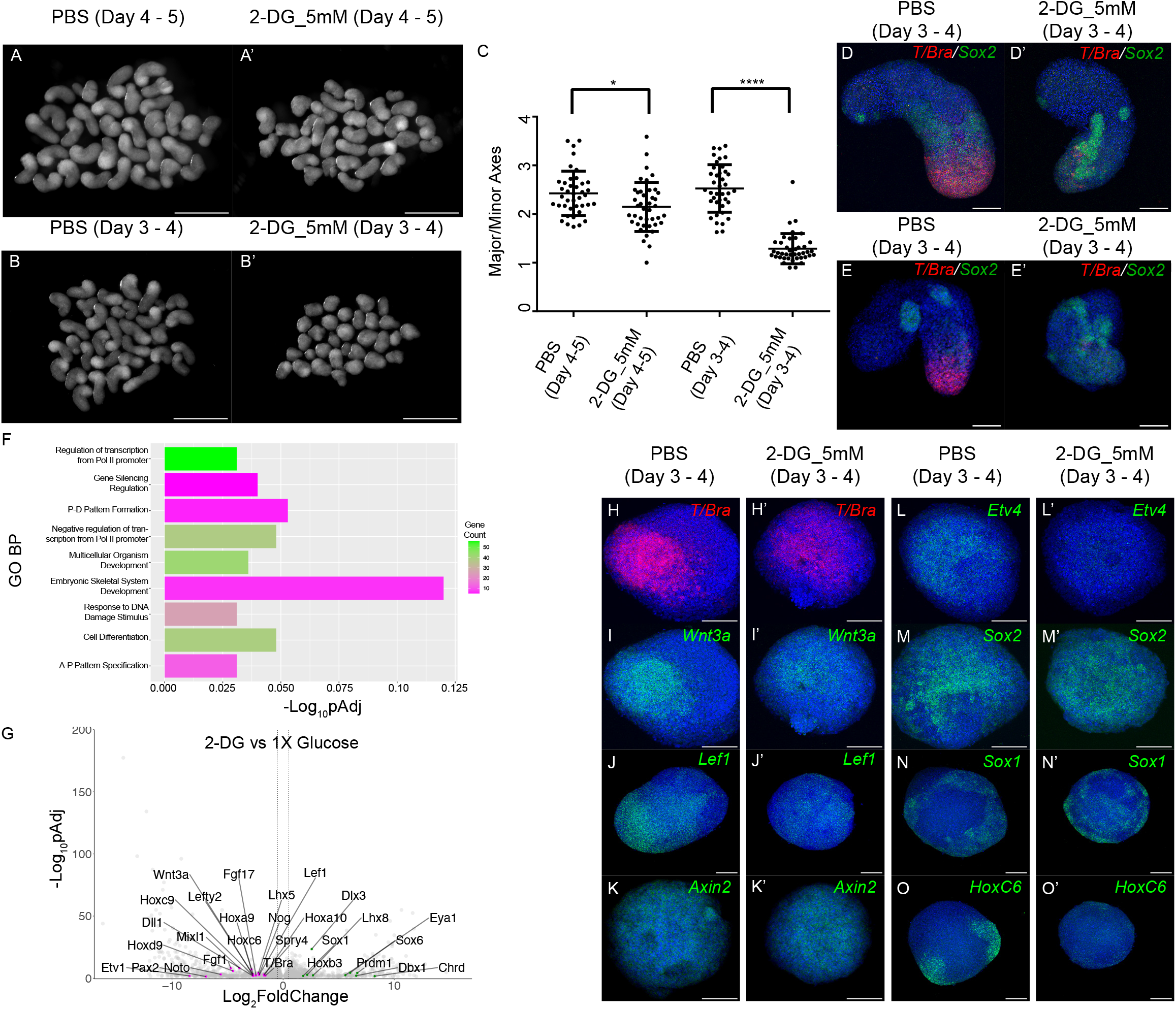
2-DG abrogates mesoderm specification. (A – A’) Overview images showing day 5 gastruloids treated with PBS (A) and 2-DG (A’) between day 4 and 5 (N=1) (B – B’) Overview images showing day 5 gastruloids treated with PBS (B) and 2-DG (B’) between day 3 and 4 (N>3) (C) Graph showing the ratio of the major axis to minor axis of the gastruloids calculated from (A-B’) images. (N =1, PBS (day 4 -5) n= 40, 2-DG (day 4 -5) n = 41, PBS (day 3 - 4) n = 39, 2- DG (day 3 - 4) n = 41). P value for PBS (day 4 – 5) vs 2-DG (day 4 – 5) = 0.0108 and P value for PBS (day 3 – 4) vs 2-DG (day 3 – 4) < 0.0001 (D-E’) Maximum intensity projections of HCR staining showing the expression pattern of *T/Bra* (Red) and *Sox2* (Green) in day 5 gastruloids treated with PBS (Day 4 – 5) (D, n=17), 2- DG (4 – 5) (D’, n=17), PBS (Day 3 – 5) (E, n=19) and 2-DG (Day 3 – 5) (E’, n=18). (F) Bar graph showing gene ontology term analysis for biological processes (GO-BP) of the downregulated genes in 2-DG compared to PBS treated gastruloids. FDR < 0.1. (G) Volcano plot showing the upregulated (Green dots) and downregulated genes (Magenta dots) in 2-DG treated gastruloids. (Cut off for Log_2_FoldChange = -1.5,1.5) (H-M’) Maximum intensity projections of HCR staining showing the expression patterns of *T/Bra* (H, n>15), *Wnt3a* (I, n=12), *Lef1* (J, n = 6), *Axin2* (K, n =12), *Etv4* (L, n =12), *Sox2* (M, n = 18), Sox1 (N, n=14) and *Hoxc6* (O, n=10) in day 4 gastruloids treated with PBS (Day 3-4) and *T/Bra* (H’, n>15), *Wnt3a* (I’, n=11), Lef1 (J’, n = 6), *Axin2* (K’, n =11), *Etv4* (L’, n =12), *Sox2* (M’, n = 16), Sox1 (N’, n=12) and *Hoxc6* (O’, n=10) in day 4 gastruloids treated with 2-DG (Day 3- 4). Scale bar – 1mm for the overview images (A-B’) and 100 μm for the confocal images (D-E’ and H-M’). The blue staining denotes nuclei in all the confocal images.

### Glucose deprivation does not impact mesoderm specification

Since 2-DG not only blocks glycolysis but has been shown to have pleiotropic effects (e.g., induction of autophagy, apoptosis, interference with protein glycosylation causing ER stress) we wanted to confirm the 2-DG mediated phenotype is linked to central glucose metabolism by completely removing glucose from the medium and assessing whether a similar phenotype is observed to that of 2-DG treatment (Pajak et al., 2019a). In addition, we also wanted to test whether higher glucose concentration led to higher expression of mesodermal markers. We incubated gastruloids in medium with no glucose (0X) or 3X higher glucose concentration between day 3 and 4 and again transferred them to the medium with 1X glucose concentration until day 5. Surprisingly, completely removing glucose from or adding 3X glucose to the medium did not lead to any elongation phenotype (Figure 3A-A’’’). The ratio of the major axis to the minor axis remained unchanged in both the absence of glucose or 3X glucose conditions (Figure 3B). This lack of elongation phenotype in 0X and 3X glucose was further confirmed with HCR analysis for *T/Bra*. 0X, 1X and 3X glucose-incubated gastruloids all showed similar expression levels of *T/Bra* (Figure 3C-C’’). To quantitatively test whether these treatments impact gastruloid growth, we compared their volumes and found a significant decrease in gastruloid volumes of 0X glucose and 2-DG treated gastruloids as compared to 1X glucose treated gastruloids (Figure 3D). Transcriptomic analysis of 0X, 1X, and 3X glucose-treated gastruloids did not enrich any class of genes which are involved in developmental processes, unlike 2-DG treatment. Nevertheless, we picked some genes which were known to have developmental roles as showed in the volcano plot (Figure 3E and 3F). We did not observe any differential expression for the genes specific to mesodermal cells. Overall, these results showed that 2-DG affects mesoderm specification independently of its glycolysis glucose inhibitor function.

**Figure 3:**
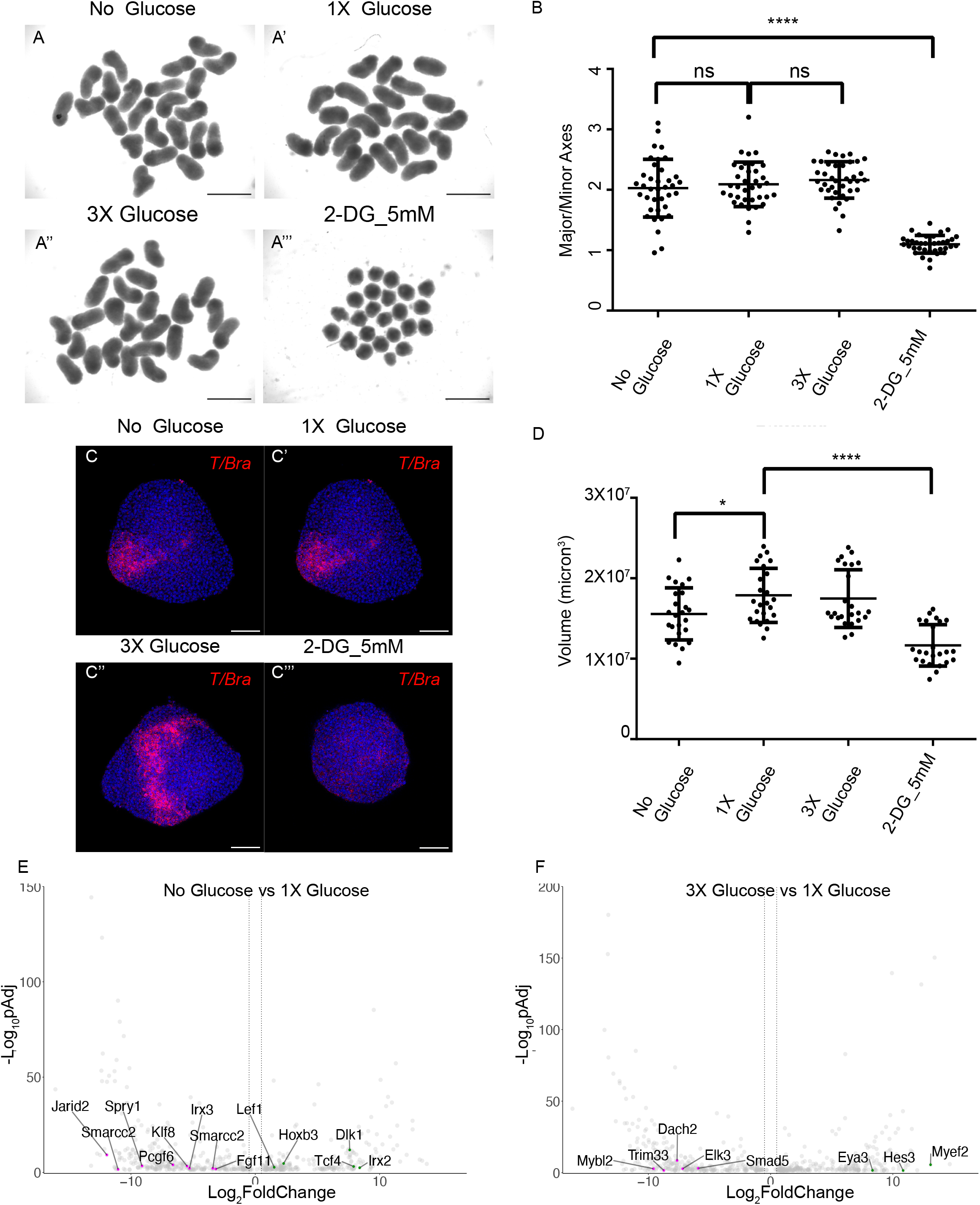
Complete removal of glucose does not affect mesoderm specification. (A – A’’’) Overview images showing day 5 gastruloids treated with 0X glucose (A), 1X glucose (A’), 3X glucose (A’’) and 2-DG (A’’’) between day 3 and 4 (N=4) (B) Graph showing the ratio of the major axis to minor axis of the gastruloids calculated from (A-A’’’) images. (N =2, 0X glucose n= 37, 1X glucose n =36, 3X glucose n =38 and 2-DG n =37). P value for 0X glucose vs 1X glucose = 0.5286, P value for 1X glucose vs 3X glucose = 0.3510, and P value 0X glucose vs 2-DG < 0.0001. (C-C’’’) Maximum intensity projections of the HCR staining showing the expression pattern of *T/Bra* in 0X glucose (C), 1X glucose (C’), 3X glucose (C’’) and 2-DG (C’’’) (N =3, n=10 each experiment). (D) Graph showing volumes of the gastruloids treated with different concentrations of glucose and 2-DG (N =3, 0X glucose, 1X glucose, and 2-DG, N=3 n =24). P value for 0X glucose vs 1X glucose = 0.0194 and P value 1X glucose vs 2-DG < 0.0001. (E) Volcano plot showing the upregulated (Green dots) and downregulated genes (Magenta dots) in 0X glucose condition compared to 1X glucose. (Cut off for Log_2_FoldChange = - 1.5,1.5) (F) Volcano plot showing the upregulated (Green dots) and downregulated genes (Magenta dots) in 3X glucose condition compared to 1X glucose. (Cut off for Log_2_FoldChange = - 1.5,1.5) Scale bar – 1mm for the overview images (A-B’) and 100 μm for the confocal images (D-E’ and H-M’). The blue staining denotes nuclei in all the confocal images.

### 2-DG treated gastruloids have a distinct metabolic signature as compared to glucose deprived gastruloids

To better understand the discrepancy between 2-DG and no glucose conditions, we needed to understand how these treatments affected glucose metabolism and other related metabolic pathways. At this end, we performed untargeted metabolomics analysis of 2-DG, 0X and 3X glucose conditions and compared this with 1X glucose. We first normalised the data by cell number to adjust metabolite levels to any differences in the sample size of the input material. A principal component analysis (PCA) showed that 2-DG treated samples are clearly distinct from the rest of the experimental conditions (Figure 4A). To further confirm that the observed difference in the 2DG treated gastruloid metabolome was not an artifact of our normalisation method, we performed another PCA independent of the cell number where we normalised the mass spectrometry (MS) signal for each metabolite detected to the sum of all the MS signals detected in each condition (Figure 4A’). This analysis also showed 2-DG treated samples to separate from rest of the experimental conditions highlighting their very distinct metabolic signature. 0X glucose and 3X glucose treated samples were still found to be closely related to the control (1X glucose) in the PCA plots (Figure 4A and A’).

**Figure 4:**
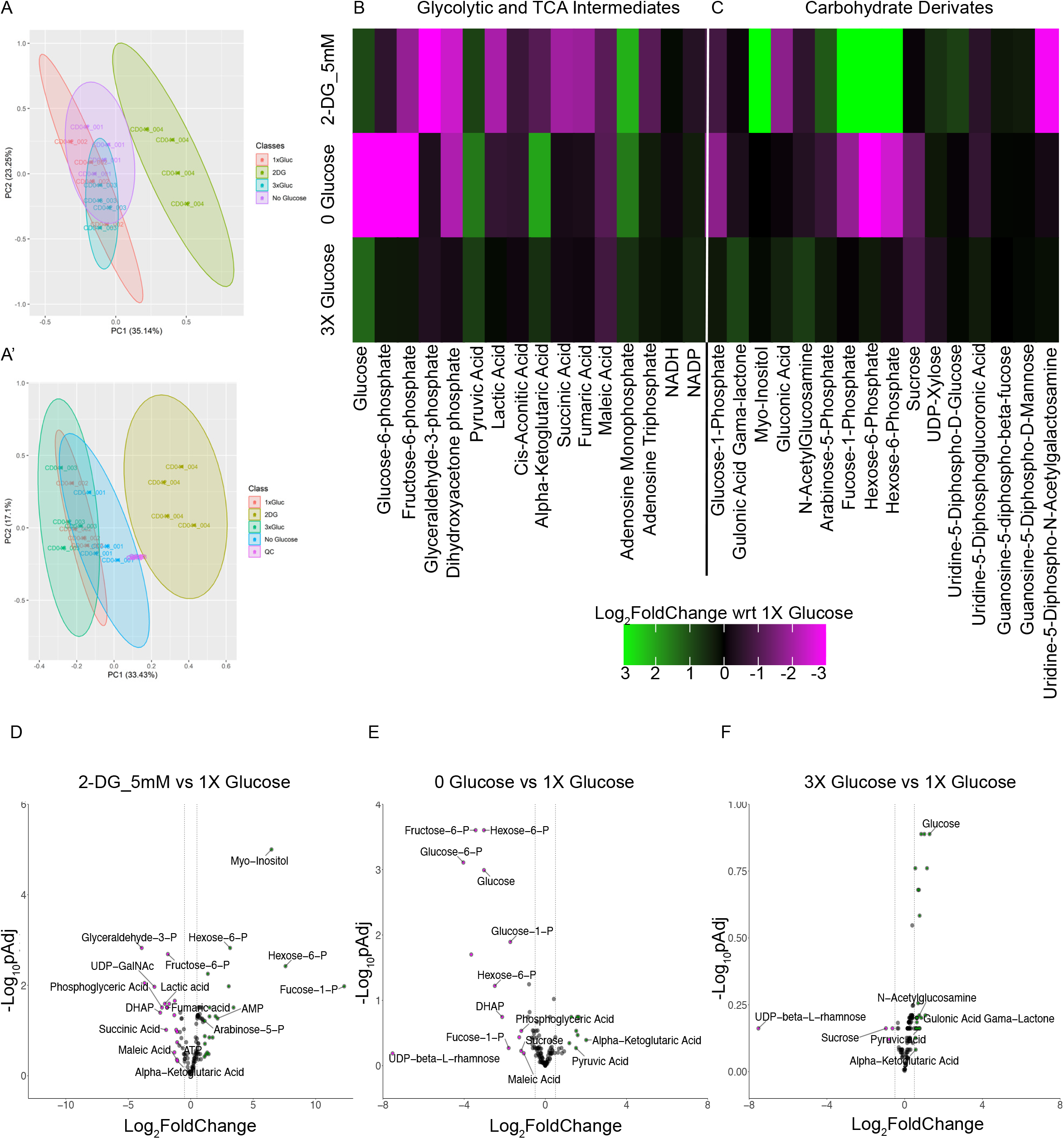
Metabolomics analysis of gastruloids grown in different glucose concentrations and treated with 2-DG. (A-A’) Variance scaled principal component analysis of 4 classes (0, 1X, 3X and 0X glucose) normalised to the total cell numbers in each class (A) and signal of each metabolite normalised to the sum of all the signals within the sample (A’). Numbers within the plot denote each class (001-004) and the biological replicate number (CD41-CD44). (B) Heatmap showing the Log_2_FoldChange in the level of glycolytic intermediates and metabolites related to carbohydrate metabolism in 0X glucose, 3X glucose and 2-DG conditions, all compared to 1X glucose. (C-E) Volcano plot showing individual metabolites in lower abundance (Magenta dots) and higher abundance (Green dots) in 2-DG (C), 0X glucose (D) and 3X glucose (E), all compared to 1X glucose. (Cut off for Log_2_FoldChange = -0.5,0.5)

At the pathway level, we found a reduction in the levels of glycolytic intermediates in the 2-DG treated condition from the level of fructose-6-P, confirming that 2-DG indeed blocks the glycolytic pathway (Figure 4B). Notable intermediates reduced in level included fructose-6-P, glyceraldehyde-3-P, and importantly lactic acid (Figure 4B and 4D). The level of ATP was mildly reduced while that of AMP was moderately increased upon 2DG treatment (Figure 4B). TCA intermediates also showed an overall decrease in their levels (Figure 4B and 4D). However, unlike in 2-DG conditions, glucose deprivation drastically affected the glycolytic pathway from the level of glucose (Figure 4B). Importantly, glucose concentration was significantly lower than that in the control confirming the efficiency of the treatment (Figure 4B and 4E). Other intermediates that were depleted included glucose-6-P, fructose-6-P, and dihydroxyacetone phosphate (Figure 4E). However, unlike in 2-DG, neither lactic acid nor TCA intermediates were impacted (Figure 4B). In addition, ATP levels were not significantly different from that in the control while AMP levels were mildly increased (Figure 4B). One possible explanation for why TCA intermediates were not impacted by the absence of glucose is that cells might take up pyruvate directly from the media in which we are culturing all gastruloids. Consistent with this idea, we see that pyruvate levels were mildly higher in the absence of glucose as compared to the control (Figure 4B and 4E).

Another important intermediate which was significantly reduced in the 0X glucose condition but unchanged in 2-DG treatment was glucose-1-P, an intermediate in the glycogen synthesis and the breakdown pathways (Figure 4B and 4E). These reduced levels could reflect either the slower production of glycogen or more rapid consumption to produce energy and other metabolic intermediates. As such, glucose-1-P could have been used to produce glucose-6-P to continue the glycolytic pathway in the 0X glucose condition.

From analysing changes in specific metabolites, we observed the expected increase in glucose levels in the 3X glucose condition compared to 1X glucose confirming the efficiency of the treatment. However, none of the glycolytic and the TCA intermediates nor any other sugar derivates showed any significant changes as compared to that in 1X glucose (Figure 4B and 4F). Nevertheless, amino acids were found to be more abundant in 3X glucose as compared to 1X glucose, hinting that excess glucose may be used in the amino acid biosynthetic pathway (Suppl. Figure 1). On the contrary, 2-DG and 0X glucose showed differences in the levels of glycolytic intermediates and sugar derivatives (Figure 4B and 4C).

This metabolomics experiment helped us to understand how different metabolic manipulations affected glycolysis both at the pathway and the individual metabolites levels. Since some of the glycolytic intermediates are funnelled into various other connected pathways, we analysed the effect of metabolic manipulations on the levels of metabolic intermediates in those pathways as well.

Glucose is used to produce other hexoses in the cell. One such sugar phosphate that was abundant in 2-DG but significantly reduced in 0X glucose was fucose-1-P (Figure 4C, 4D and 4E). Fucose is produced from glucose and is an important sugar used in post-translational modification (PTM) in its activated form GDP-fucose (Becker & Lowe, 2003). GDP-fucose itself is produced from Fucose-1-P in a reversible manner. Surprisingly, the levels of GDP-fucose were not different between 2-DG and 0X glucose treatments (Figure 4C). One explanation for this outcome in the 2-DG treated gastruloids is that fucose-1-P is also produced in the lysosome from the degradation of fucosylated proteins (Becker & Lowe, 2003). A higher level of fucose-1-P, in 2-DG conditions could therefore be a product of the degradation of fucosylated proteins as 2-DG is known to induce ER stress, unfolded protein response pathway and autophagy (Pajak et al., 2019b). In the case of no glucose conditions, the lack of glucose itself would be expected to lead to a lower level of fucose-1-P and so we had to assume that it has been used primarily to maintain the steady levels of GDP-fucose.

Our metabolomics data could detect other carbohydrate derivatives involved in glycosylation such as GDP-mannose, UDP-glucose, N-acetylglucosamine (GlcNAc) but their levels were similar in both 2-DG and 0X glucose (Figure 4C). We could not detect the activated form of GlcNAc (UDP-GlcNAc), but the level of its epimer UDP-N-AcetylGalactosamine (UDP-GalNAc) was significantly reduced in 2-DG but not upon modulating glucose levels (Figure 4C and 4D). Altogether our metabolomics dataset revealed both similarities and differences between 2-DG and 0X glucose conditions with only 2-DG impacting intermediates involved in post-translational modification.

### Mannose rescues 2-DG treatment phenotypes in mesodermal cells

We next sought to determine what the mechanism of 2-DG action is that that blocks mesoderm specification, given that this is not phenocopied by an inhibition of central glucose metabolism in response to glucose deprivation. 2-DG not only blocks the glycolytic pathway but, due its structural similarity, competes with mannose and gets incorporated into the glycosyl chains of glycosylated proteins (Figure 5A). In a recent glycoproteomic study, clusters of proteins involved in germ layer and mesoderm specification were detected to be affacted an unbiased manner upon 2-DG treatment which otherwise were mannosylated (Ma et al., 2022). This prompted us to test whether 2-DG acts independently of the glycolytic pathway.

**Figure 5:**
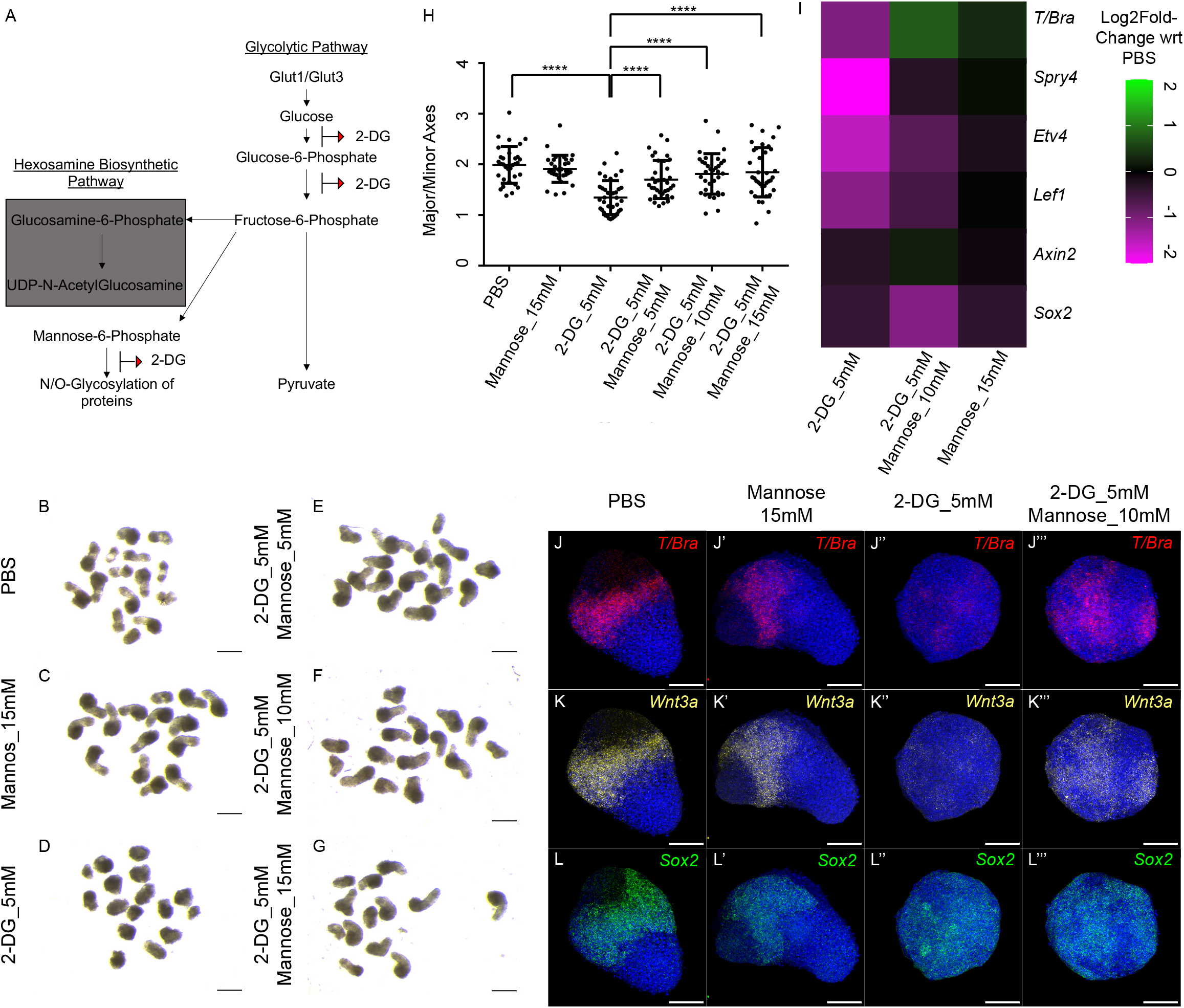
Mannose rescues the 2-DG mediated phenotype. (A) Schematic showing the glycolytic pathway and the targets of 2-DG (B-G) Overview images showing day 5 gastruloids treated with PBS (B), Mannose_15mM (C), 2-DG_5mM (D), 2-DG_5mM Mannose_5mM (E), 2-DG_5mM Mannose_10mM (F), 2-DG_5mM Mannose_15mM (G). (N = 4). (H)Graph showing the ratio of the major axis to minor axis of the gastruloids calculated from (B - G) images. (N =3, PBS n= 32, Mannose_15mM n =33, 2-DG_5mM n =37, 2-DG_5mM Mannose_5mM n =35, 2-DG_5mM Mannose_10mM n=34, 2-DG_5mM Mannose_15 mM n=36). P value for PBS vs 2-DG_5mM < 0.0001, P value for 2-DG_5mM vs 2-DG_5mM Mannose 5mM < 0.0001, P value for 2-DG_5mM vs 2-DG_5mM Mannose 10mM < 0.0001, P value for 2-DG_5mM vs 2-DG_5mM Mannose 15mM < 0.0001. (I) Heatmap showing the Log_2_FoldChange in the gene expression in 2-DG_5mM, 2-DG_5mM Mannose_10mM and Mannose_15mM compared with PBS treated gastruloids. (N=2) (J-L’’’) Maximum intensity projections of the HCR staining showing the expression patterns of *T/Bra* (J-J’’’), *Wnt3a* (K-K’’’) and *Sox2* (L-L’’’) in PBS (J,K,L n=5), Mannose_15mM (J’,K’,L’, n=8), 2-DG_5mM (J’’, K’’, L’’, n=10) and 2-DG_5mM Mannose_15mM (J’’’, K’’’, L’’’, n =10). Scale bar – 1mm for the overview images (B-G) and 100 μm for the confocal images (J-L’’’). The blue staining denotes nuclei in all the confocal images.

To determine whether mannose could rescue the phenotype by competing with 2-DG, we treated gastruloids with 5mM of 2-DG and three different concentrations of mannose (5mM, 10 mM and 15mM) between day 3 and 4. We then transferred them to medium containing no mannose and 2-DG until day 5. The overview images show that mannose alone did not affect gastruloid development as compared to the PBS control (Figure 5B and 5C). 2-DG treatment affected elongation when compared with both PBS and mannose treatments (compare Figure 5D with 5B and 5C) but when 2-DG was supplemented with mannose, the elongation was restored (compare Figure 5D with 5E, 5F and 5G), similar to the PBS control (compare Figure 5B and 5C with 5E, 5F and 5G). This rescue was also reflected in the ratio of the major axis to the minor axis of the gastruloids (Figure 5H).

To confirm whether this elongation phenotype also mirrored the restoration of mesodermal cell markers and signalling pathways, we performed qPCR analysis for *T/Bra*, FGF target genes *Etv4* and *Spry4*, the neural marker *Sox2* and the general Wnt targets *Axin2* and Lef1. As shown before, the expression of *T/Bra* was downregulated in 2-DG as compared to the PBS control gastruloids (Figure 5I). However, upon addition of 10mM mannose, the expression was restored to that in the control (Figure 5I). Alone, 15mM mannose did not have any effect on *T/Bra* expression (Figure 5I). A similar trend was observed for an Fgf target gene *Spry4* (Figure 5I). Expression levels of *Etv4* and Wnt target gene *Lef1* were reduced in 2-DG, but mannose supplementation had a very little effect on the rescue of their expression levels (Figure 5I). Neither 2-DG, 2-DG with mannose nor mannose alone led to any changes in the expression of the neural gene *Sox2* and the broad range Wnt target gene *Axin2* (Figure 5I). We further confirmed the qPCR data with HCR. Expression of *T/Bra* and *Wnt3a* was lost upon 2-DG treatment (Figure 5J’’ and 5K’’) in comparison to that in PBS control (Figure 5J and 5K). 15mM mannose has no effect on the expression of both *Wnt3a* or *T/Bra* (Figure 5J’ and 5K’). However, the expression of both *Wnt3a* and *T/Bra* can be rescued by the addition of 10mM mannose together with 2-DG (Figure 5J’’’ and 5K’’’). Expression of *Sox2* remained unchanged in all treatments (Figure 5L-L’’’). Altogether our data shows that mannose addition is sufficient to restore the expression of mesodermal genes and the morphogenetic process of axial elongation in 2-DG treated gastruloids. These results also highlight that glycosylation events (but not the glycolytic pathway) are important for the mesodermal fate specification and subsequently axial elongation.

To further confirm the importance of glycosylation in mesoderm specification, we treated gastruloids with 5mM glucosamine. Although glucosamine is important in glycosylation, in excess, it interferes with the addition of mannose and glucose residues to the growing glycosyl chains (Beriault et al., 2017). Like 2-DG treatment, glucosamine treated gastruloids appeared less elongated as compared to their control counterparts (Suppl. Figure 2A-A’). The ratio of the major to the minor axis was significantly reduced in glucosamine treated gastruloids (Suppl. Figure 2B). We further checked whether the shortening of the major axis was a result of abrogated mesodermal development. HCR staining for *T/Bra* revealed that indeed its expression was reduced in the glucosamine treated gastruloids both in day 4 and 5 gastruloids (compare Suppl. Figure 2C with 2E and 2D with 2F). Although unlike in 2-DG, the expression of *Wnt3a* was not completely diminished in day 4 gastruloids (Suppl. Figure 2C’ and 2’), in day 5 gastruloids, its expression was not localised to the posterior pole in the treated gastruloids as compared to the control (Suppl. Figure 2D’ and 2F’). Expression of *Sox2* remained unchanged (compare Suppl. Figure 2C’’with 2E’’ and 2D’’ with 2F’’). Altogether our 2-DG and glucosamine dataset highlights the importance of mannose in protein glycosylation and subsequently mesoderm specification.

## Discussion

Through an interrogation of the impact of 2-DG on gastruloid development, and comparing this to glucose deprivation, we have discovered a novel role of mannose in early mesoderm specification. In addition to blocking glycolysis, 2-DG interferes with glycosylation, specifically mannosylation, of proteins (Ma et al., 2022). Both our mannose rescue and glucosamine experiments showed that glycosylation is required for mesoderm specification. Glycosylation is an important post-translational modification that has a variety of cellular functions both in development and disease (Czajewski & Aalten, 2023; Haltiwanger & Lowe, 2004; Reily et al., 2019). The WNT and FGF signalling pathways are involved in mesoderm specification and constitute the proteins whose glycosylation and proper intracellular trafficking are important in mediating the pathway activity. Fgfr1, an FGF receptor expressed in the primitive streak, is N-glycosylated and this glycosylation is important for the regulation of the Fgf pathway (Duchesne et al., 2006; Polanska et al., 2009; Zukowska et al., 2023). N-glycosylation of *Wnt3a*, which is specifically expressed in the mesodermal cells, is important for its secretion and hence the activation of Wnt signalling (Komekado et al., 2007). Here we have observed a disruption in the activity of the FGF signalling pathway as a result of 2DG treatment, that can also be rescued by the addition of mannose. This provides a useful experimental system to further explore the mechanistic links between mannose metabolism and developmental signalling pathways during germ layer patterning. These findings are in line with recent work in the mouse embryo showing that disruption of late-stage glycolysis does not disrupt primitive streak elongation and mesoderm specification, which instead depends on the modulation of FGF signalling via glycosylation (Cao et al., 2023).

Having established the role of mannose in mesoderm specification, the important question is what the sources of mannose are, since gastruloids are cultured in the medium containing only glucose. Under physiological conditions, one mannose source comes from glycolysis in which fructose-6-P gets converted into mannose-6-P and later mannose-1-P, which is used in glycosylation through its activated form GDP-Mannose (Ichikawa et al., 2014; Sharma et al., 2014). This still calls into question how the intracellular levels of mannose are maintained when glucose is completely removed from the medium, as our metabolomics data showed no change the levels of GDP-Mannose in the 0X glucose condition as compared to the control (Figure 4B). Under such conditions, previous studies have shown that glycogen reserves serve as a source of mannose which is later used up during protein glycosylation. An additional source could be the glycoproteins from which it gets recycled (Fujita et al., 2008; Ichikawa et al., 2014). Under such limited supply of mannose, when we treated 0X glucose gastruloids with 2-DG we observed a much more severe phenotype (Suppl. Figure 3A-A’’). Metabolic tracer experiments would be useful to elucidate how mannose levels are regulated in mesodermal cells and how metabolic remodelling takes places to restore the mannose reserves under 0X glucose condition.

This study contributes to a recent body of work focussed on exploring links between central carbon metabolism and body plan formation in vertebrate embryos (Bulusu et al., 2017; Miyazawa et al., 2022; Oginuma et al., 2017, 2020). A previous study in the chicken embryo showed that 2-DG blocked mesoderm specification in the tailbud in a glycolysis dependent manner, which subsequently increased lactic acid secretion and increased intracellular pH (Oginuma et al., 2017, 2020). While we have observed that mannose treatment is sufficient to rescue the 2-DG inhibition of mesoderm markers in the context of gastruloid symmetry breaking, we cannot rule out that an additional regulation of Wnt signalling via modifying intracellular pH is an additional mechanistic link between glycolysis and mesoderm specification in this context. Indeed, this could be the case in gastruloids as lactic acid levels were lower in 2-DG treated gastruloids as compared to the control (Figure 4B).

Despite not requiring glucose during the onset of gastruloid elongation, a strong expression of glucose transporters co-localises with *Sox2* and *T/Bra* expression during these stages (Figure 1), raising the question of their function in both of these cell types. One hypothesis is that these cells do not require glucose in the specification process but rather in other cellular processes that occur downstream and are not captured in the gastruloid assay. During gastrulation, mesodermal cells undergo an EMT (Dias et al., 2020; Ferrer-Vaquer et al., 2010; Guibentif et al., 2021; Morgani & Hadjantonakis, 2019), which is an energy expensive process (Kang et al., 2019; Sciacovelli & Frezza, 2017). A previous study using the mouse embryo showed how culturing PSM explants in glucose-free medium impacted its development and segmentation (Bulusu et al., 2017; Miyazawa et al., 2022). Indeed, recent work in the mouse embryo has demonstrated that late-stage glycolysis becomes important only after ingression through the primitive streak (Cao et al., 2023). Together with this work, this reveals an emerging picture of how differential hexose metabolism can be used to drive successive stages of gastrulation through the modulation of key signalling pathways such as FGF.

## Acknowledgments

We acknowledge the support of the EMBL Metabolomics Core Facility (MCF) in the acquisition and analysis of liquid chromatography-mass spectrometry data. We would like to additionally thank Bernhard Drotleff for advice on sample preparation for these experiments. BJS was supported by a Henry Dale Fellowship jointly funded by the Wellcome Trust and the Royal Society (109408/Z/15/Z), and a Wellcome Trust Discovery Award (225360_Z_22_Z). This work was funded by MRC Research Grant MR/V009192/1 jointly held by CD and BJS. We thank Lara Busby and Alexandra Neaverson for critically reading the manuscript, Kevin Costello for the bioinformatics analysis and Tamsin Samuels for help with the R script and graphs.

## Materials and Methods

### Culturing and maintenance of mouse embryonic stem cells

E14 cells were bought from ATCC (CRL-1821, ATCC) and cultured on the mouse embryonic fibroblasts (MEFs) for the first 3 passages. Later, the cells were weaned off the MEFs and cultured again for 2-3 passages before cry-preserving them storing them. For the initial 5-6 passages, cells were cultured in the GMEM (11710035, ThermoFisher Scientific) along with 10% ES grade foetal bovine serum (FBS) (16141002, ThermoFisher Scientific), 1000 units/ml Leukaemia Inhibitory Factor (LIF) (ESG1106, Merck), 100 uM Beta-mercaptoethanol (31350010, ThermoFisher Scientific), 1X Non-Essential amino acids (11140050, ThermoFisher Scientific), 1mM of Sodium Pyruvate (11360070, ThermoFisher Scientific) and 2mM L-Glutamine (25030149, ThermoFisher Scientific). At this stage, cells were frozen in this medium supplemented with 10% DMSO (D2438, Merck). To plate gastruloids, frozen cells were thawed and cultured for 3 passages in 2iLIF medium which contained a Wnt agonist, CHIR99021 (3uM, SML1046, Merck), ERK antagonist PD 0325901 (1uM, PZ0162-5MG, Merck) and 1000 units/ml LIF diluted in N2B27. N2B27 medium was made using the N2 (0.5X) and B27 (0.5X) (17504044, ThermoFisher Scientific) supplements in 1:1 DMEM/F12 (21331020, ThermoFisher Scientific) to Neurobasal media (21103049, ThermoFisher Scientific). N2 supplement was prepared in house as described previously (Mulas et al., 2019). Cells were routinely checked for mycoplasma.

### Culturing Gastruloids in different glucose concentrations, inhibitor, and mannose treatments

200-300 ES cells per gastruloid were plated and cultured as previously described by Baillie-Johnson et al. (Baillie-Johnson et al., 2015). For the 2-DG (D8375, Merck) treatment, 500mM stock solution was made in PBS (P4474, Merck) and filter sterilised using a 0.22 um filter. Gastruloids were treated between day 3 and 4 or day 4 and 5 with 5mM 2-DG diluted in N2B27. For differential glucose concentration treatments between day 2 and 3 and between day 3 and 4, N2B27 was made using DMEM/F12 without glucose (L0091-500, Biowest) and Neurobasal without glucose (A2477501, ThermoFisher). For 1X and 3X glucose concentrations, 1.1 M glucose stock solution was used (A2494001, ThermoFisher Scientific) to obtain the final concentrations of 21.25mM and 63.75mM respectively. Final pyruvate concentration was adjusted to 0.36mM by adding sodium pyruvate. For the mannose (M6020, Merck) and glucosamine (G1514-100G, Merck) treatment, 500mM stock was prepared in PBS and filter sterilised. 5, 10 and 15mM working solutions were made in N2B27 with or without 5mM 2-DG. These treatments were performed on gastruloids grown in the same plate. One plate was kept as a control until day 5 to ensure that control gastruloids in the same plate elongated properly, otherwise the entire plate was discarded.

### Fixing, HCR staining, immunostaining and imaging of the gastruloids

Day 4 and day 5 gastruloids were harvested from 96 well plates, washed in PBS and fixed overnight in 4% paraformaldehyde (158127, Merck). The next day, gastruloids were dehydrated in a series of increasing methanol concentrations and stored at -20 in 100% methanol at least for a day before using them for the HCR experiments. Briefly, the HCR protocol involved rehydrating the gastruloids with a descending series of methanol concentrations and finally PBS-T (0.1% Tween, 655204, Merck). Once hydrated, gastruloids were incubated in the pre-hybridisation (pre-hyb) buffer at 37 degrees for 30 minutes and then the pre-hyb buffer was replaced with HCR probes diluted in the pre-hyb buffer. Version 3.0 HCR Gene specific probes were designed by Molecular Instruments. Hybridisation took place through incubating the gastruloids with probe solution overnight at 37 degrees. The next day, the probe solution was replaced and gastruloids were washed 4 times with the washing buffer for 15 minutes at 37 degrees. This was followed by 3 × 5 minute washes with 5X Sodium Chloride Sodium Citrate + 0.1% Tween-20 (5X SSC-T) solution. Gastruloids were then incubated with the Hairpin Amplification Buffer for 30 minutes. The fluorophore-tethered hairpins were briefly denatured before diluting them in the amplification buffer. Gastruloids were incubated overnight in the dark with the hairpin solution at room temperature. The next day, gastruloids were washed with 5X SSC-T and then PBS-T before staining them with DAPI overnight (D1306, ThermoFisherScientific). Finally, gastruloids were cleared for 24 hours in ScaleS4 (Hama et al., 2015) and imaged on the confocal microscope. All the HCR reagents and probes were bought from Molecular Instruments Inc, USA.

For the whole-mount antibody staining, gastruloids were fixed in 4% PFA overnight and washed with PBS-T three times on a shaker for 5 mins per wash. Later, the blocking, primary, and secondary antibody incubations and washes were performed as described previously with the exception of performing all the steps in a 2 ml tube (Baillie-Johnson et al., 2015). DAPI was added to the secondary antibody solution. After completing the washes, gastruloids were cleared with ScaleS4 as described above. The list of antibodies is provided in the supplementary section.

HCR or antibody stained gastruloids were imaged by mounting them in ScaleS4 sandwiched between two coverslips on the Zeiss LSM700 with Plan-Apochromat 20X objective.

### Metabolomics analysis

Day 4 gastruloids from 2 plates were collected (approximately 45-48 gastruloids, per treatment, per biological replicate) (N = 4) into ice cold PBS in a clean glass petri dish, washed once and transferred to another glass petri dish on ice containing cold PBS. Gastruloids were collected, briefly centrifuged to settle them down and the PBS was completely removed. Gastruloids were snap-frozen in liquid nitrogen. To count the number of cells in each gastruloid, 5 gastruloids from each batch and each treatment, were collected, trypsinised for 1 min in 50 ul of 0.05% trypsin (25300054, ThermoFisher Scientific) and neutralised using 950 ul of warm N2B27. Cells were immediately counted using the Neubauer haemocytometer chamber. Snap-frozen gastruloids were sent to EMBL-Heidelberg, Germany, for the metabolomics analysis.

### Transcriptomics analysis

Day 4 gastruloids from 1 plate (approximately 20-24 gastruloids, per treatment, per biological replicate) (N, biological replicate = 3) were collected in ice cold PBS, excess PBS was removed from the tube and initially 100 ul of TRIzol (15596026, ThermoFisher Scientific) was added to dissociate and homogenise the gastruloids. Later, 900 ul of TRIzol was added and stored in -80 until RNA extraction and library preparation. RNA was extracted using the TRIzol protocol and diluted in nuclease-free water. RNA concentration was measured using the QuBit RNA High Sensitivity kit (Q32852, ThermoFisher Scientific). For the library preparation, mRNA was isolated from 2.5 ug of total RNA using NEBnext Poly (A) magnetic isolation module (E7490S, NEB). The library was prepared as per the user’s manual using the NEBNext Ultra II Directional RNA library Prep Kit for Illumina (E7760S, NEB). Library indexing was done with NEBNex multiplex oligos for Illumina Set 1 (E7335S, NEB) and Set 2 (E7500S, NEB). Quality of the final library preparation was analysed on the Bioanalyzer using the High Sensitivity DNA reagents kit and cassettes (5067-4626, Agilent). For all the steps involving purification of library intermediates, SPRISelect magnetic beads (B23317, Beckman Coulter) were used. The transcriptomics experiment itself was outsourced to the Cancer Research UK, Cambridge. Reads obtained were analysed using the online tool, Galaxy. The pipeline consisted of trimming the index sequences using Trim_galore, obtaining BAM/SAM files using HISAT2 (Kim et al., 2019), removing PCR duplicates using Picard Mark Duplicates followed by StringTie to assemble transcripts (Pertea et al., 2015) and DESeq2 for obtaining differentially expressed genes (Love et al., 2014). Gene ontology analysis was performed using the online tool DAVID.

### qPCR analysis of candidate genes

RNA was extracted as described above. cDNA was prepared using the Superscript III first-strand cDNA synthesis kit (18080051, ThermoFisher Scientific). For the standardisation of primers, control cDNA was diluted as 1:40, 1:80, 1:160 and 1:320 and the Ct values were calculated for each primer pair for a gene and compared with that of the internal control H2az1 (Veazey & Golding, 2011). Standardised primer sets were then used to calculate the gene expression change in different treatment groups. For this experiment, 1:80 cDNA dilution was used. A list of the primers used is provided in the supplementary material. Statistical Analysis and graphs

For statistical analysis and graphs shown in Figure, 2C, 3B, 3D, 5H and Supp Figure 2B, GraphPad Prism was used. For volcano plots shown in Figure 2G, 3E, 3F, 4D, 4E and 4F, for the heatmaps shown in Figure 4B, 4C, 5I, Supp Figure 1 and the bar graph shown in Figure 2G, the R package ggplot2 was used.

## Supplementary Figure legends

Supp. Figure 1: Metabolomics analysis of amino acids, nucleotides, and fatty acids in gastruloids treated different glucose concentration and 2-DG.

(A-C) Heatmap showing the Log_2_FoldChange in the level of amino acids (A), nucleotides (B) and fatty acids (C) in 2-DG_5mM, 0X glucose and 3X glucose, all compared to 1X glucose.

Supp. Figure 2: Excess glucosamine blocks mesoderm specification and axial elongation.

(A-A’) Overview images showing day 5 gastruloids treated with PBS (A) and 5mM glucosamine (A’) between day 3-4.

(B) Graph showing the ratio of the major axis to minor axis of the gastruloids calculated from (A-A’) images. (N =2, PBS n= 28, Glucosamine_5mM n =30) P value for PBS vs Glucosamine_5mM < 0.0001

(C-F’’) Maximum intensity projections of HCR stains showing the expression patterns of *T/Bra* (C, D, E, F), *Wnt3a* (C’, D’, E’, F’) and *Sox2* (C’’, D’’, E’’, F’’) in PBS (C-D’’, day 4 n=9 and day 5 n =6) and Glucosamine_5mM (E-F’’, day 4 n=10 and day 5 n = 7)

Scale bar – 1mm for the overview images (A-A’) and 100 μm for the confocal images (D-E’’). The blue staining denotes nuclei in all the confocal images.

Supp. Figure 3: 2-DG treatment in 0X glucose condition leads to much severe condition.

(A-A’’) Overview images showing day 5 gastruloids treated with 1X glucose (A), 1X glucose with 5mM 2-DG (A’) and 0X glucose with 5mM 2-DG (A’’) between day 3-4.

Scale bar – 1mm for the overview images

## Supplementary Material and Methods

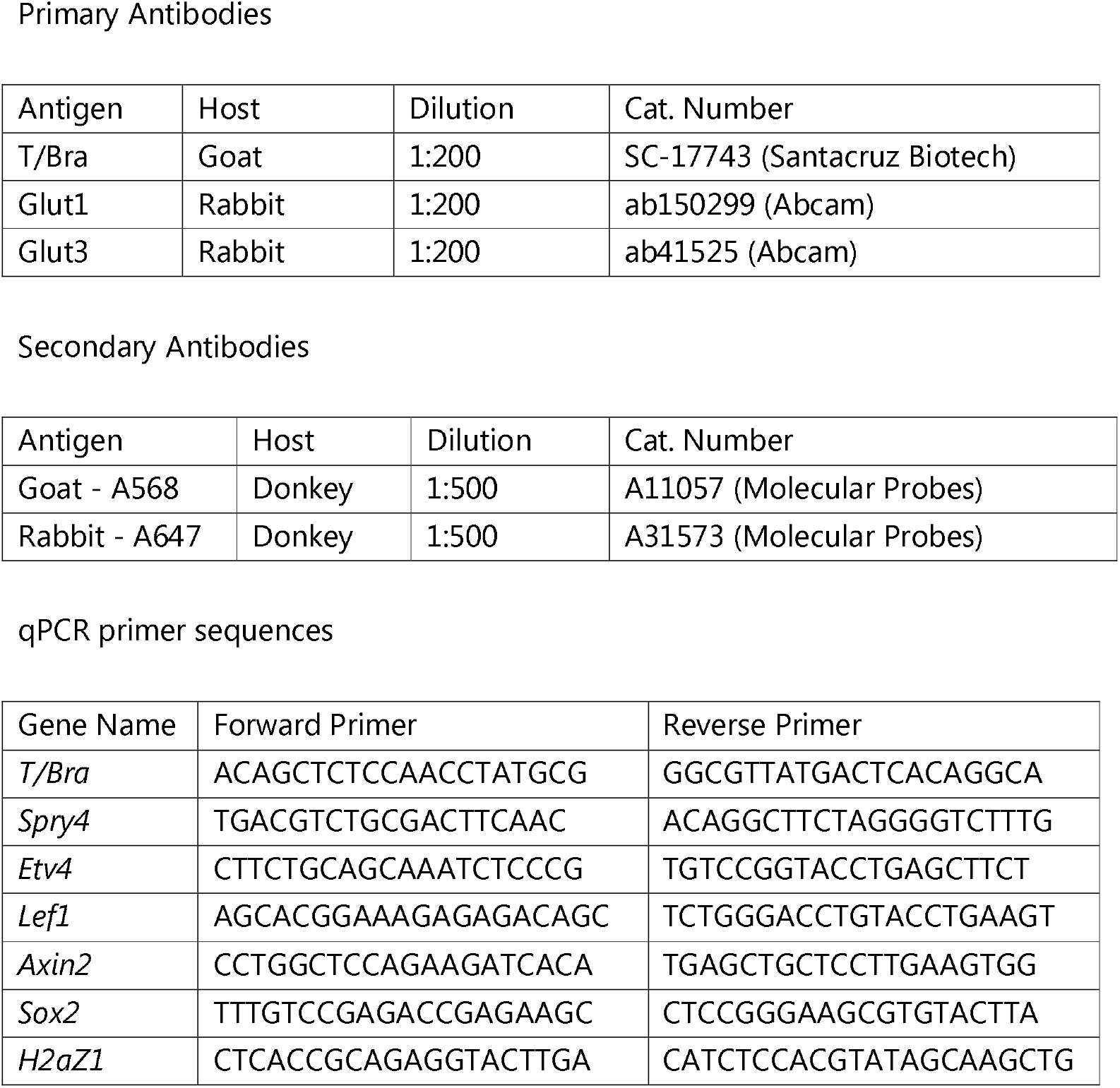

