## Supplementary figures and images for "Mannose is crucial for mesoderm specification and symmetry breaking in gastruloids"

### Supplementary Figure

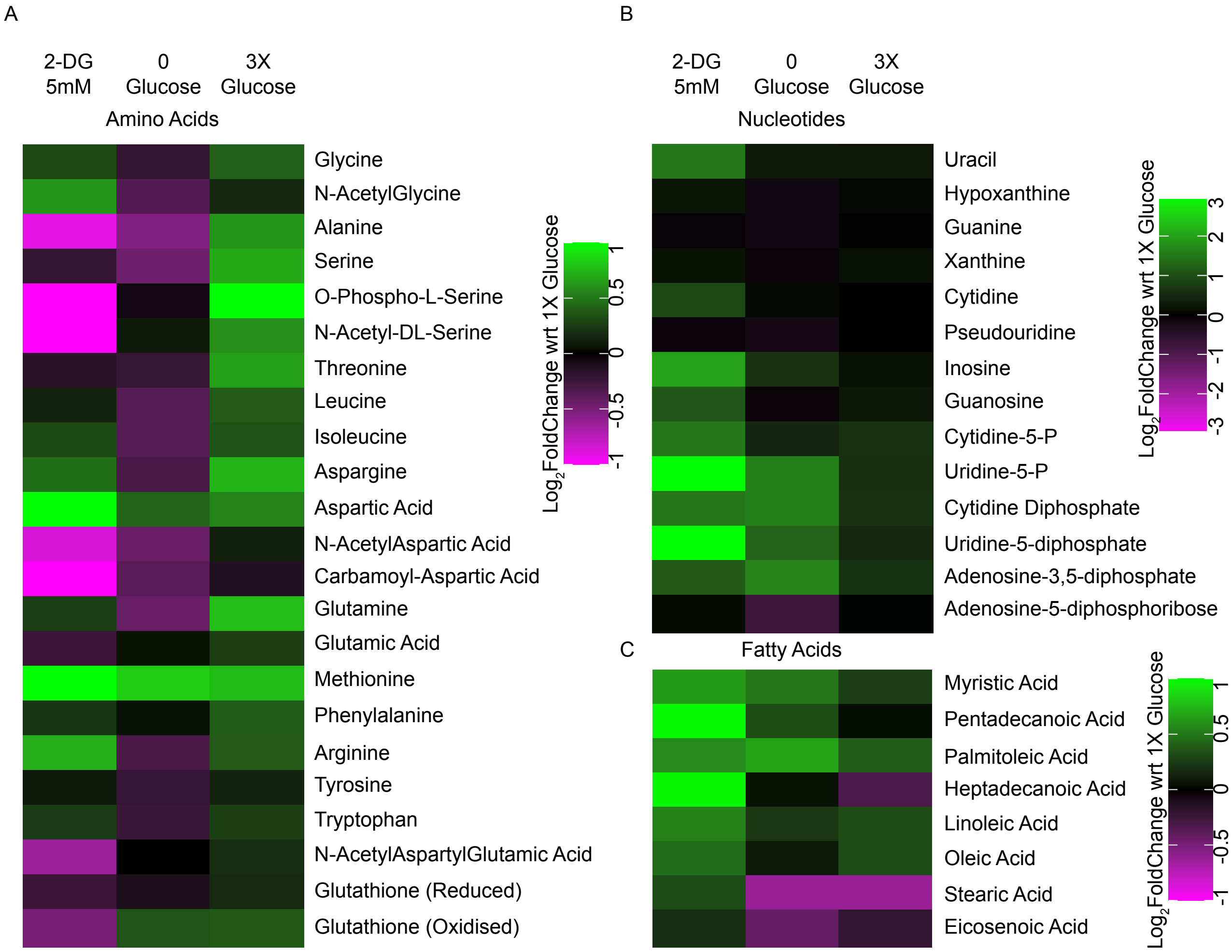

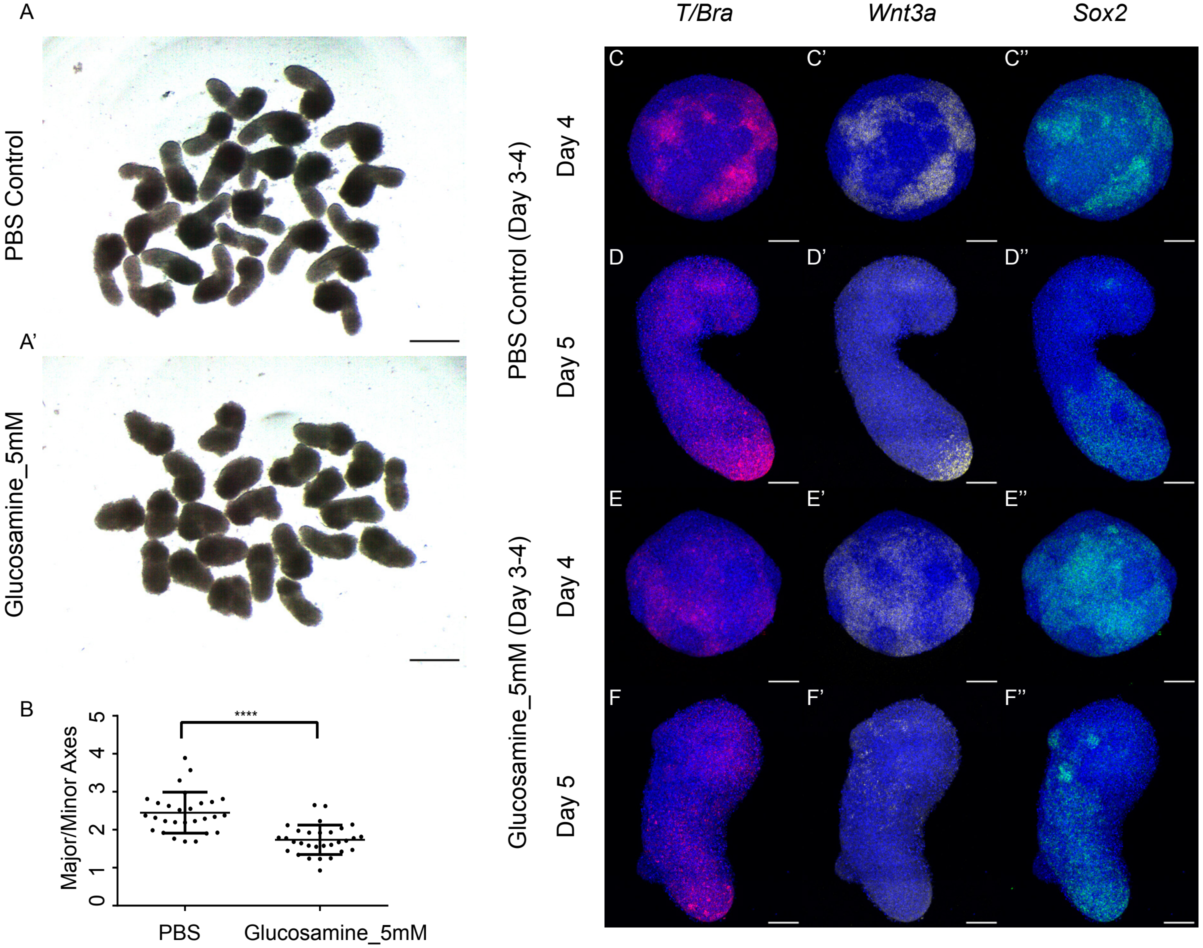

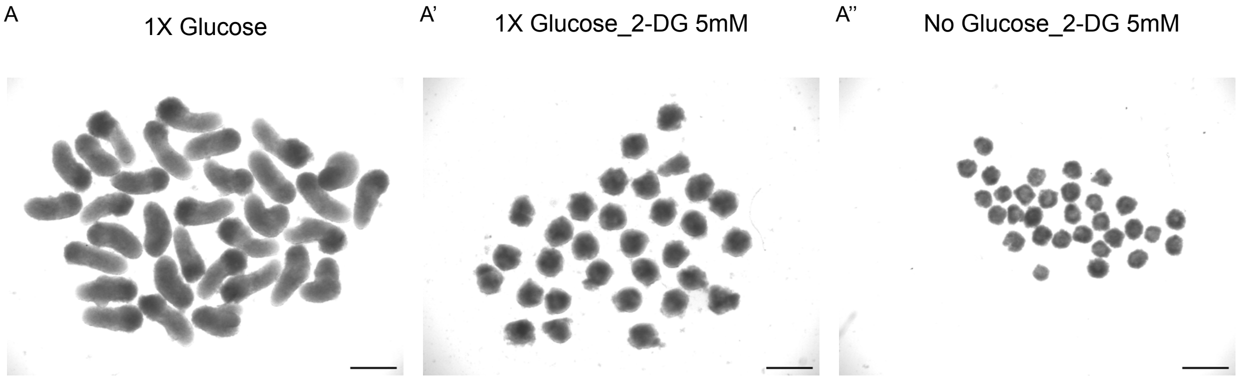
